# Calibration-free compression brings Evo 2 to its full million-token context on a single GPU

**DOI:** 10.64898/2026.08.28.747902

**Authors:** Michail Patsakis, Alexandros Tzanakakis, Ilias Georgakopoulos-Soares

## Abstract

Evo 2 is the largest openly available genomic foundation model, but its forty billion parameter configuration cannot be loaded onto a single 80 GB accelerator, placing genome-scale analysis beyond most laboratories. We present TurboQuant-Bio, an open toolkit that compresses Evo 2’s weights and attention cache to four bits without calibration data, and serves both through fused kernels. Compression is near-lossless across perplexity spanning the tree of life, genomic classification, splice-site prediction, gene completion and clinically relevant variant-effect prediction. It brings Evo 2 40B onto one 80 GB GPU and Evo 2 7B to its full million-token context within a 40 GB memory budget, an eightfold gain in reachable context. We further show that the released chunked-prefill path is silently incorrect, returning plausible but uncorrelated likelihoods, and derive the block-wise continuation that repairs it: a complete 580-kilobase bacterial genome is now scored in one context in 22 minutes rather than 13.7 hours.

## Introduction

Evo 2 is currently the largest openly available genomic foundation model [1]. Built on the StripedHyena-2 hybrid architecture [2,3] and trained on trillions of nucleotides drawn from every domain of life, it predicts the effects of genetic variation from regulatory elements to clinically important genes, and it generates genome-scale sequence, all without task-specific fine-tuning. It extends an earlier line of long-context genomic models [4], and complements supervised long-range models of regulatory activity such as Enformer [16] and is designed to read context windows of up to one million bases, which is what lets it place a local sequence in its broader genomic context. For a researcher choosing a single model to build on, Evo 2 is among the most capable options available, which is why making it easy to run matters.

Capability of this kind is expensive to deploy, and the memory Evo 2 consumes at inference has two parts that are easy to conflate but behave very differently. The first is the fixed cost of the parameters, paid once when the model is loaded, which dominates when the context is short. The second is the cost of the attention cache, the running record of past positions the model consults as it reads a sequence. This second cost grows with sequence length and is well known to dominate long-context inference [5,6,7], so it comes to dominate in the long-context regime. In full precision the forty-billion-parameter configuration does not fit on a single graphics processor, and its long-context memory can exceed even a well-provisioned multi-GPU node. The seven-billion-parameter configuration fits, but only until the context grows, at which point it too runs out of memory. The result is a hardware barrier that leaves one of the most powerful open genomic foundation models, and its most distinctive capability, out of reach for much of the research community.

Compression is the natural way to lower this barrier, and post-training quantization, which stores a trained model at lower numerical precision, is the most direct form of it. The methods that reach the best fidelity depend on calibration data, a representative sample of inputs used to tune the compression to the model, whether through error correction, salient-channel protection or fitted rotations [8,9,10,11]. In biology this dependence is a real obstacle. High-quality in-domain data are often scarce, and Evo 2 is routinely applied across organisms and assays whose statistics differ enough that no one calibration set is representative. A method that must be re-tuned whenever the input distribution shifts is awkward to use and hard to trust. What is needed instead is compression that requires no calibration, so that it can be applied to a new Evo 2 model release immediately and reused unchanged across the whole tree of life.

Recent work has shown that calibration-free quantization need not sacrifice fidelity. TurboQuant [12] uses a geometric idea: a random rotation applied to a high-dimensional vector makes the distribution of its coordinates predictable in advance, after which a fixed quantizer, designed once for that predictable distribution, can be applied with no knowledge of the data. PolarQuant [13] shares the calibration-free, geometry-first premise but reaches it differently, preconditioning the vector and then quantizing in polar coordinates rather than coordinate by coordinate. This gives a principled route to compression without calibration, related in spirit to incoherence-based weight quantization [14]. It has been developed and demonstrated primarily for the attention caches of conventional transformer models, and adapted recently to the attention caches of protein language models [15]. Calibration-free integer schemes for attention caches are likewise well established for conventional decoders [6,7]. Whether any of this transfers to the weights and to the unusual hybrid architecture of Evo 2, whether it preserves the biological behaviour Evo 2 is valued for, and whether the resulting system is fast enough to be practical, has not been established.

Here we answer these questions for Evo 2 and, in doing so, remove two distinct obstacles to long-context genomic modelling. We present TurboQuant-Bio, an open toolkit that applies calibration-free compression jointly to the weights and the attention cache of Evo 2, together with fused kernels that serve the compressed model efficiently. The first obstacle is hardware. With no calibration data and only small changes in biological performance, Evo 2 40B runs on a single 80 GB GPU, where the uncompressed model cannot even be loaded, and long-context inference with Evo 2 7B fits within a 40 GB memory budget, the capacity of accelerators many laboratories already have. The second obstacle is correctness, and it is one we did not expect to find. Beyond the length of a single forward pass, long sequences must be processed in chunks; we show that the released chunked implementation silently returns wrong likelihoods, and we derive the block-wise continuation that repairs it. Compression is exposed as two deployment tiers, one that reduces the growing memory of the attention cache while keeping generation as fast as full precision, and one that additionally compresses the parameters for the smallest possible footprint, so that a user can match the trade-off to the hardware in front of them. Below we show that biological behaviour is preserved across perplexity spanning the tree of life, zero-shot genomic classification and splice-site prediction, clinically relevant variant-effect prediction and gene completion; that the repaired long-context path is faithful and fast; and that the whole system runs on one GPU.

## Results

### Weight memory compression

We first quantified how much calibration-free weight quantization reduces Evo 2’s fixed memory footprint, which must be paid before any sequence is processed and determines how much capacity remains for long-context activations and cache storage. For the linear layers that are quantized, stored weight memory decreases from 60.8 to 15.7 GiB. Total on-device allocated memory after model loading decreases from 76.6 to 31.7 GiB (**Fig. 1A**). The on-device reduction is smaller than the storage reduction because the per-block scales and the small set of unquantized parameters are kept in full precision. In practical terms the full-precision model leaves almost no free memory on a single H100 GPU, whereas the compressed model leaves tens of gigabytes of headroom for the activations and cache that long context requires.

**Figure 1.**
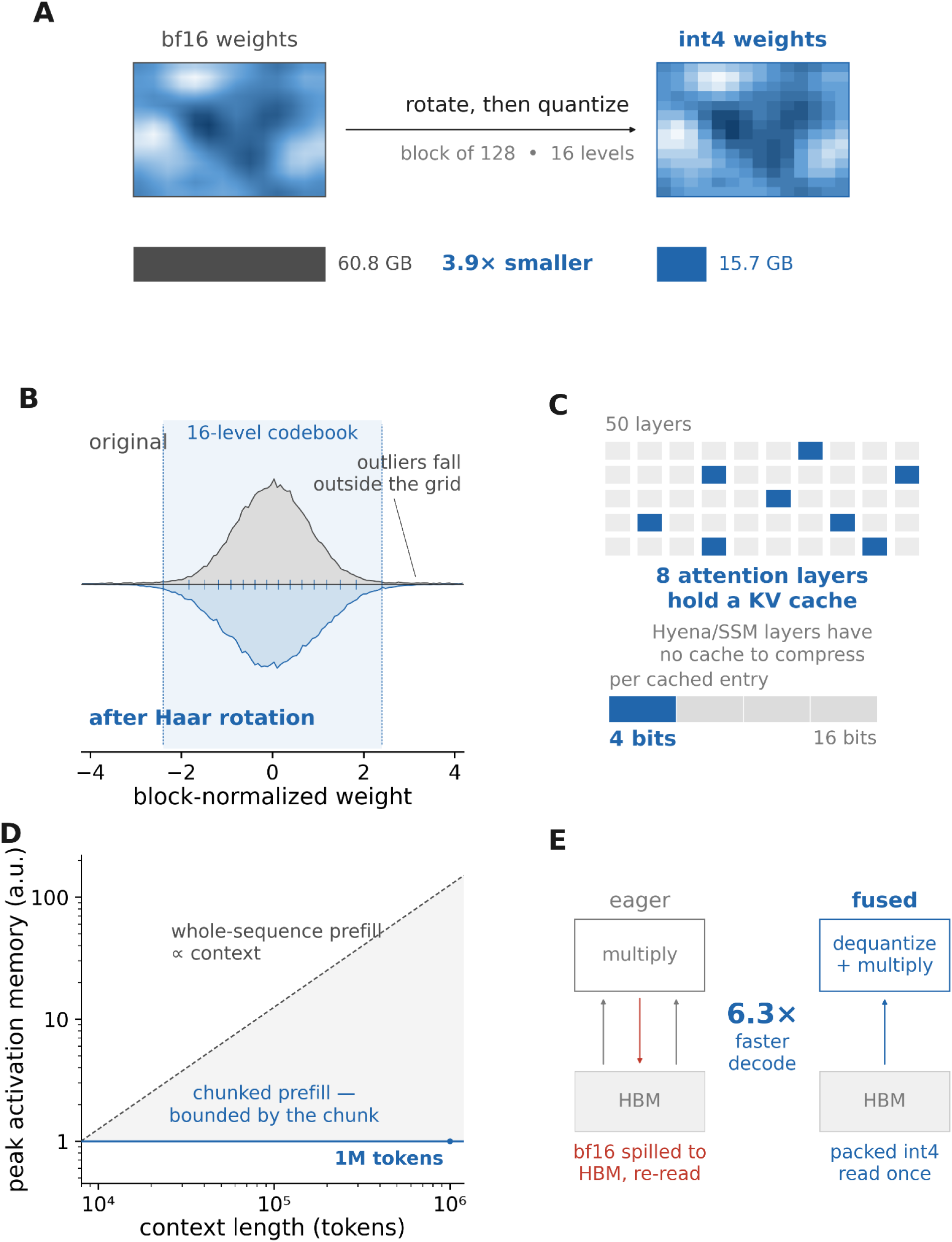
Four-bit compression of Evo 2 40B. **(A)** Weight blocks are rotated and quantized to 16 levels, reducing the storage of the quantized linear-layer weights from 60.8 to 15.7 GiB **(B)** The Haar rotation removes the heavy tail, so one fixed 16-level codebook (shaded band, ticks at the Lloyd–Max levels) covers every block. **(C)** Only 8 of 50 layers hold a KV cache; each entry is stored in four bits rather than sixteen. **(D)** Chunked prefill bounds peak activation memory by the chunk rather than the context, making 10⁶-token inputs tractable. **(E)** Fused kernels dequantize in registers, so packed int4 weights and K/V are read from HBM once, giving 6.3× faster decode than the eager streaming path. Panels a, b, d and e are schematic.

### Perplexity fidelity across bit-widths and the tree of life

Next, we examined to what extent Evo 2 could be quantized without compromising its language-model fidelity [1]. Four-bit quantization is near-lossless for language-model fidelity. On the T4 phage genome, the geometric-mean perplexity of Evo 2 40B rises by 0.51% at four-bit. The effect is small in magnitude but consistent across all ten windows (10/10 increases; sign test p = 0.002), while eight-bit is unchanged within rounding and two-bit degrades substantially (**Fig. 2B**). This places four-bit as the natural operating point, delivering substantial compression while leaving the model’s fidelity essentially unchanged.

**Figure 2.**
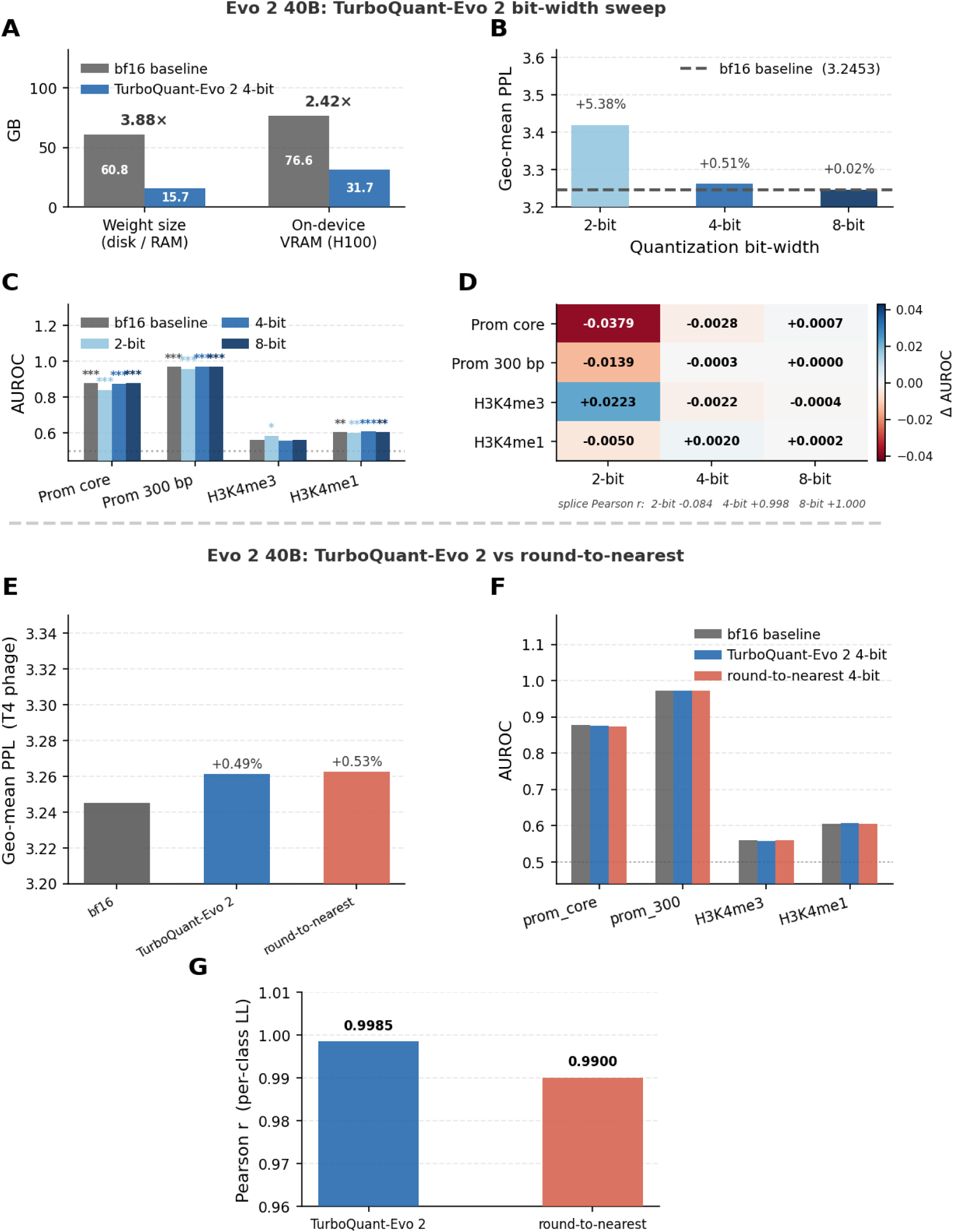
Four-bit weight quantization preserves accuracy on Evo 2 40B, and TurboQuant-Bio compares favorably with round-to-nearest. **(A)** Weight storage and on-device memory, full precision versus four-bit. **(B)** Geometric-mean perplexity on T4 phage windows by bit-width. **(C)** Zero-shot GUE classification AUROC, all conditions besides H3K4me3, significantly above chance. **(D)** Change in AUROC and splice-site correlation by bit-width. Eight-bit is effectively lossless, four-bit is near-lossless, two-bit is too aggressive for fine-grained discrimination. **(E)** Full precision, TurboQuant-Bio four-bit and round-to-nearest four-bit compared on T4-phage perplexity. **(F)** GUE classification AUROC **(G)** and splice-site correlation.

Because Evo 2 is trained on all domains of life [1], a compression method is only useful if it is faithful across that diversity rather than on a single genome. We measured four-bit perplexity on fourteen taxa spanning viruses, bacteria, archaea and eukaryotes, and compared TurboQuant-Bio against round-to-nearest at the same bit-width (**Supplementary Table 1**). Baseline perplexity varies widely across taxa, reflecting the composition of the training distribution. Against that backdrop the four-bit degradation stays below two percent for every taxon and is largest for the most GC-rich and high-diversity genomes. Across all fourteen taxa, TurboQuant-Bio produces a smaller or more favorable perplexity change than round-to-nearest, and its advantage is greatest exactly where quantization is hardest, which is the behaviour a calibration-free method should have if its fidelity comes from principle rather than from a fitted distribution.

### Zero-shot downstream tasks

We next asked whether the fidelity preserved at four-bit extends from sequence modeling to downstream biological tasks. Overall, we find that compression preserves the model’s downstream biological behaviour. On the four GUE genomic classification tasks [17], four-bit AUROC changes by no more than a few thousandths and none of the changes are statistically significant (**Fig. 2C**, **D**). Splice-site prediction, a more fine-grained task, retains near-perfect correlation with full precision at four-bit and eight-bit but collapses at two-bit, a clear demonstration that two-bit is too aggressive for fine discrimination even when coarse classification survives. Against the calibration-free round-to-nearest baseline, TurboQuant-Bio is closer to full precision on cross-taxa perplexity, where it produces a smaller or more favourable perplexity change on all fourteen taxa (**Supplementary Table 1**). On the single-genome T4 measurement and on the four GUE tasks the two calibration-free schemes are statistically indistinguishable at this sample size, differing by 0.02 percentage points of perplexity and by at most 0.005 AUROC in either direction (Fig. 2E-G). The separation between them emerges only across the diversity of the tree of life, which is where a calibration-free method is under the most pressure. Together, these results identify four-bit quantization as the best trade-off between compression and preservation of downstream biological performance.

### Cache compression enables million-token inference on Evo 2 40B

Weight compression reduces the fixed cost of the parameters, but the memory that ultimately limits long-context inference is the attention cache, which grows with sequence length. To separate and then combine the two effects, we measured peak and resident memory for Evo 2 40B on a single four-GPU node across context lengths from thirty-two thousand to one million tokens, for four configurations: the uncompressed baseline, four-bit weights alone, four-bit cache alone, and both together (**Fig. 3**).

**Figure 3.**
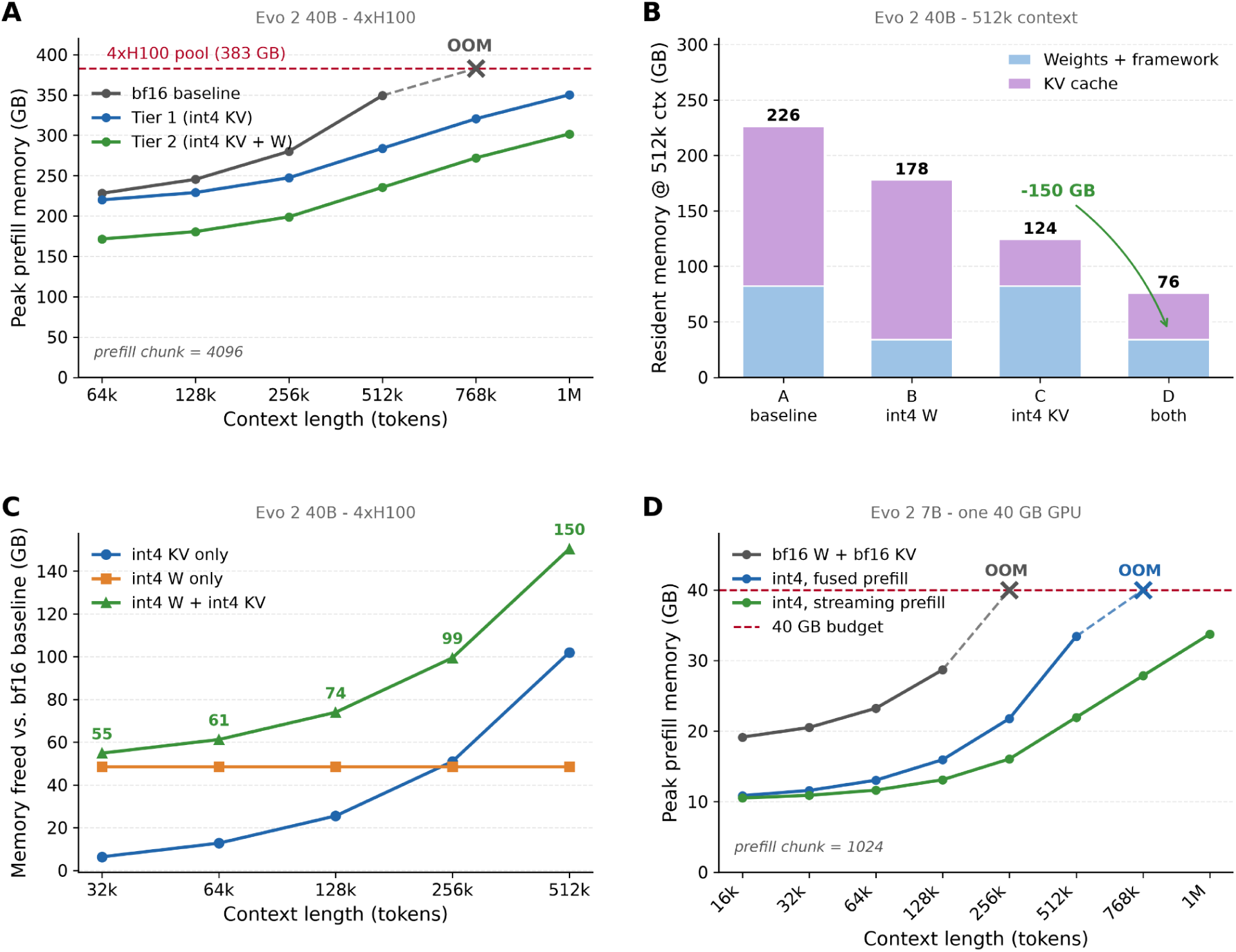
Evo 2 40B on 4× H100 unless noted. **(A)** Peak prefill memory versus context at prefill chunk 4096; the uncompressed baseline runs out of memory beyond 524,288 tokens while both compressed tiers reach one million. **(B)** Resident memory at 524,288 tokens — the longest context at which all four configurations run — split into weights and cache, prefill chunk 2048. **(C)** Memory freed relative to the uncompressed baseline for four-bit cache, four-bit weights, and both, prefill chunk 2048. **(D)** Evo 2 7B peak prefill memory on a single H100 constrained to a 40 GB memory budget, prefill chunk 1024. Full precision fails beyond 131,072 tokens; the compressed model reaches 524,288 under the default fused prefill and the full 1,048,576 when the cache is streamed one block at a time during the prompt pass.

The uncompressed baseline’s peak memory rises steeply with context and exhausts the four-GPU pool beyond 524,288 tokens, while both compressed tiers reach one million (**Fig. 3A**). Decomposing resident memory at 524,288 tokens — the longest context at which all four configurations still run — separates the two effects: the cache accounts for 144 GB against 82 GB of weights, so it is both the larger term and the one that grows (**Fig. 3B**). The difference in how the two compressions scale is the central systems result (**Fig. 3C**). Because the cache grows with sequence length while the weights do not, the memory freed by cache compression grows with context, whereas the memory freed by weight compression is constant at roughly 48 GB. Below about 256,000 tokens weight compression frees more memory; above it, cache compression dominates and is what keeps the model within budget. Combining both frees the most at every context and, at 524,288 tokens, reduces resident memory from 226 GB to 76 GB (Fig. 3B); at one million tokens the compressed model needs 175.0 GB of resident memory and peaks at 301.7 GB, where full precision cannot run at all (**Table 1**). To our knowledge this is the first demonstration that Evo 2 40B fits, in memory, at its full one-million-token context on a single node, enabled specifically by compressing the cache. We report this as a memory result; the fidelity of long-context inference is a separate question, and one the next section shows had to be repaired before it could be answered.

**Table 1.** Deployment envelope for Evo 2 under four-bit compression. Maximum context is the longest sequence scored without exhausting memory. Peak and resident memory are summed over devices for the 40B and measured on the single device for the 7B. Prefill throughput is the context length divided by the time to score it, measured at each row’s maximum context, so rows at different maxima are not directly comparable. Prefill chunk 1024 for the 7B and the single-GPU 40B, 4096 for the multi-GPU 40B. The 7B rows use an H100 constrained to a 40 GB budget; the 7B million-token row uses streaming prefill (with the default fused prefill the maximum is 524,288). Resident memory was not recorded in the 7B sweep. The 40B in full precision cannot be loaded onto one 80 GB device at all, its weights alone requiring 82.3 GB.

| Model | Hardware | Configuration | Max context | Resident (GB) | Peak (GB) |
| --- | --- | --- | --- | --- | --- |
| 7B | 1 GPU, 40 GB budget | bf16 W + bf16 KV | 131,072 | — | 28.7 |
| 7B | 1 GPU, 40 GB budget | int4 W + int4 KV | 1,048,576 | — | 33.7 |
| 40B | 1 × H100 80 GB | bf16 W + bf16 KV | cannot load | — | — |
| 40B | 1 × H100 80 GB | int4 W + int4 KV | 131,072 | 33.8 | 62.8 |
| 40B | 4 × H100 | bf16 W + bf16 KV | 524,288 | 231.4 | 349.3 |
| 40B | 4 × H100 | bf16 W + int4 KV | 1,000,000 | 223.4 | 350.2 |
| 40B | 4 × H100 | int4 W + int4 KV | 1,000,000 | 175.0 | 301.7 |

### Memory controls, cache fidelity, and fused-kernel performance

Beyond the choice of what to compress, peak memory is governed by two largely independent controls (**Fig. 4A-C**). The first is the size of the chunk used to process a long prompt, which sets the transient convolutional-transform buffer; smaller chunks lower the prompt peak at the cost of slower prompt processing, and this control is orthogonal to cache compression, which shrinks the persistent cache. The second is the cache bit-width, which trades fidelity for memory (**Fig. 4B**). Relative to full precision, eight-bit cache is near-lossless but frees the least memory, two-bit is substantially lossy, and four-bit is the operating point that preserves fidelity while freeing a large share of the cache. Streaming attention is essential to realize these savings; in long context it nearly halves the peak memory of generation compared with reconstructing the whole cache at once, at identical resident footprint, because it reconstructs one block at a time (**Fig. 4C**).

**Figure 4.**
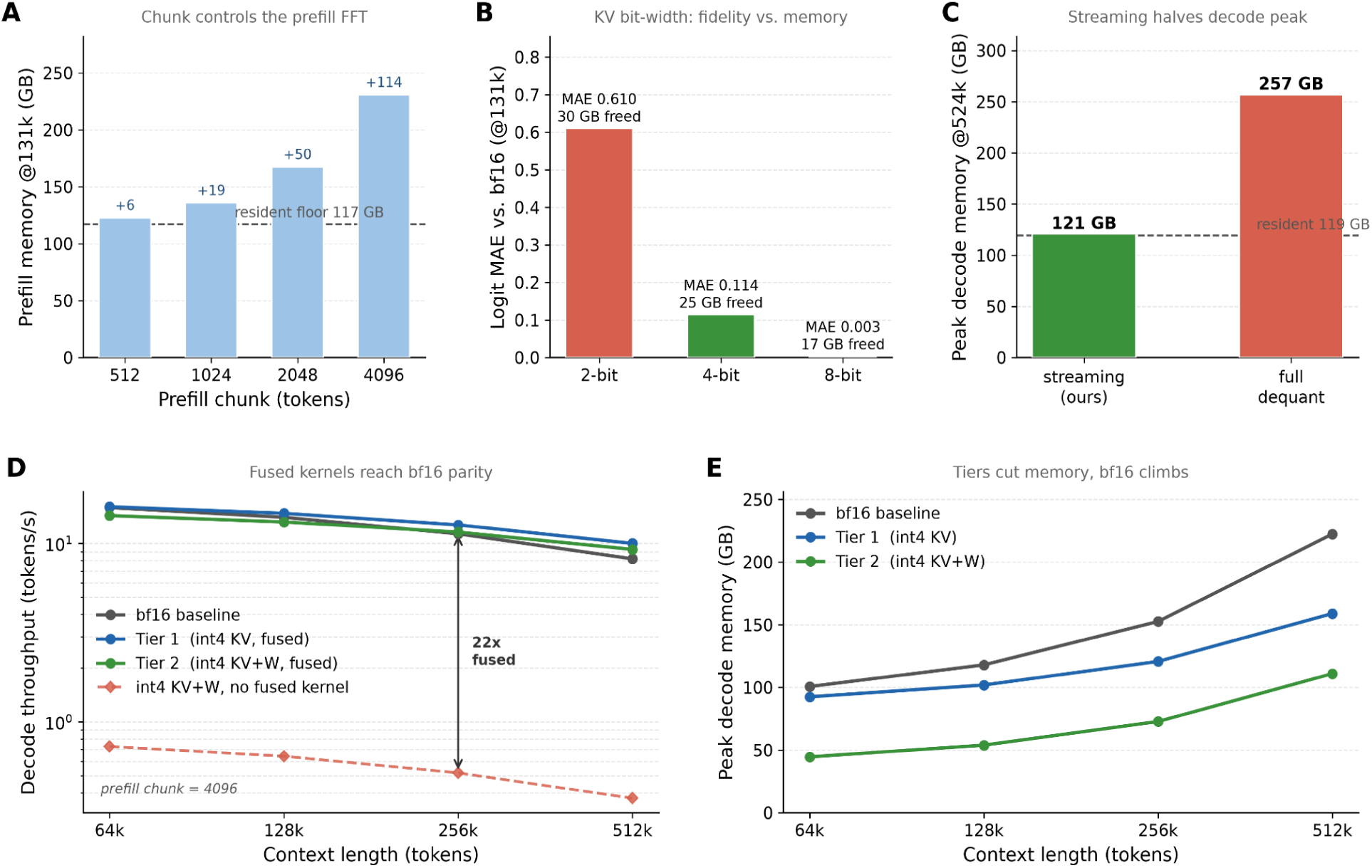
Memory mechanisms and fused-kernel performance for Evo 2 40B (4x H100). Top row, memory and fidelity mechanisms: **(A)** prompt peak memory and prompt speed versus chunk size, showing the chunk controls the transient prefill buffer; **(B)** fidelity versus cache bit-width and memory freed; **(C)** peak decode memory, streaming versus reconstructing the whole cache. Bottom row, fused-kernel results: **(D)** decode throughput versus context for full precision, Tier 1 (int4 cache) and Tier 2 (int4 cache and weights), all with the fused kernels, plus the same int4 configuration without a fused kernel; the fused tiers sit at full-precision parity while the unfused path is roughly an order of magnitude slower. **(E)** Peak decode memory versus context; the tiers cut memory substantially while full precision climbs.

The memory savings must not come at a lasting cost to speed, and with the fused kernels described in Methods they do not (**Fig. 4D, E**). With cache compression alone (Tier 1), generation throughput matches full precision at the contexts where full precision still runs, and is faster at long context, because the operation the kernel accelerates grows with context while the uncompressed model’s memory ceiling arrives first. Beyond the point where the uncompressed model runs out of memory, the compressed model sustains competitive throughput all the way to one million tokens, a context the uncompressed model cannot reach, while using substantially less memory at every shared context. Adding weight compression (Tier 2), the fused weight kernel keeps generation within 14 percent at the shortest context measured, narrowing as context grows, matching full precision by 262,144 tokens and exceeding it at the longest shared context. In short, the smallest-footprint tier is now also fast enough to use for generation, not only for memory-bound batch analysis. The fused kernels are applied transparently: the same generate call that uses the packed cache invokes them automatically, so practitioners obtain the speed benefit without any change to their code. We conclude that these optimizations convert compression into a practical long-context regime, reducing peak memory while preserving generation throughput at contexts that full precision cannot reach.

### Variant-effect prediction is robust to model compression

To establish that compression remains suitable for biologically meaningful inference, we tested whether it preserves Evo 2’s ability to score functional genetic variants. Compression preserves performance on this clinically relevant benchmark. We evaluated Evo 2 40B on the BRCA1 saturation-genome-editing assay of Findlay and colleagues [18], scoring each single-nucleotide variant by its zero-shot log-likelihood difference from the reference over a genomic window of several kilobases. On eight hundred variants, of which 165 are annotated loss-of-function and 635 functional or intermediate, the full-precision model separates loss-of-function from the pooled functional and intermediate classes with an AUROC of 0.908 and correlates with the experimental function score with a Spearman coefficient of 0.551. Applying four-bit weight quantization, with the attention cache retained in full precision, leaves these essentially unchanged, at an AUROC of 0.906 and a Spearman coefficient of 0.528, retaining almost all of the full-precision performance (**Fig. 5A, B, Supplementary Table 2**). At the level of individual variants the compressed and full-precision scores are nearly identical, with a per-variant correlation of 0.992, indicating that four-bit quantization introduces only small, unbiased perturbations to the likelihood landscape rather than degrading its variant-effect signal. The roughly fourfold weight-memory reduction is therefore obtained with negligible loss in a task with direct clinical relevance.

**Figure 5.**
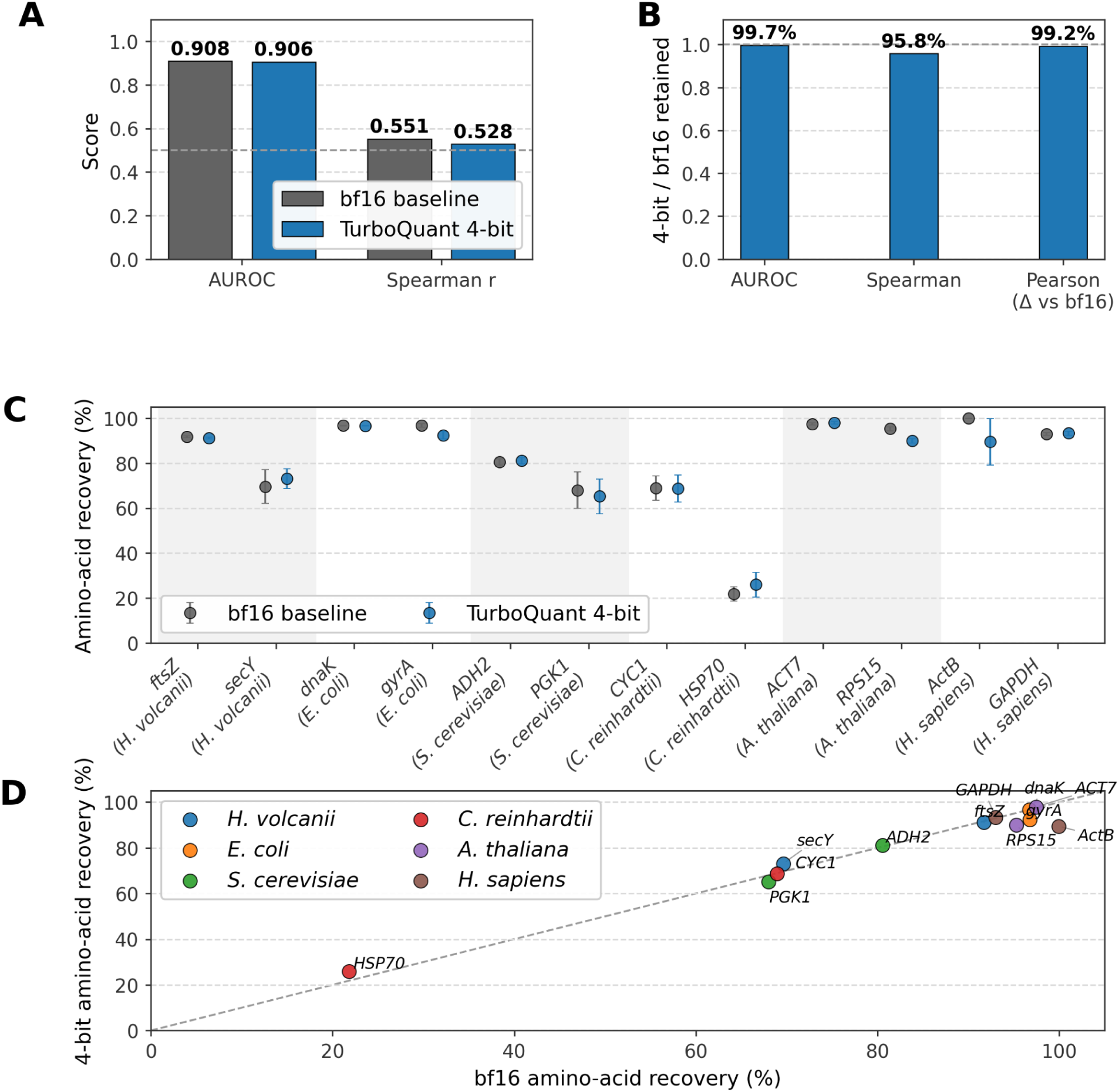
Effect of TurboQuant 4-bit quantization on Evo 2-40B variant-effect prediction and gene completion. **(A)** Variant-effect prediction performance (AUROC and Spearman correlation) for the full-precision (bf16) and 4-bit models. **(B)** Agreement between the two models: fraction of bf16 performance retained at 4-bit and the per-variant correlation of their scores. **(C)** Per-gene amino-acid recovery of the generated sequence for bf16 and 4-bit, across the 12-gene Evo 2 gene-completion panel (4 prokaryotic/archaeal genes: *ftsZ*, *secY*, *dnaK*, *gyrA*; 8 eukaryotic genes: *PGK1*, *ADH2*, *CYC1*, *HSP70*, *RPS15*, *ACT7*, *ActB*, *GAPDH*). Points show the mean over 8 stochastic generations per gene; error bars, SEM. **(D)** The corresponding per-gene mean amino-acid recovery plotted bf16 against 4-bit, colored by organism; dashed line indicates equality.

### TurboQuant-Bio preserves Evo 2 gene-completion performance

To test whether compression harms Evo 2’s generative capability, we benchmarked the 40B model on the Evo 2 gene-completion panel of twelve genes spanning six organisms from archaea to human [1], replicating the gene set and scoring methodology of the original Evo 2 study. For each gene, the model was given an upstream context and a coding-sequence prefix and asked to generate the remainder of the gene, after which amino-acid recovery was scored over the held-out region. The bf16 and compressed conditions used the same eight random seeds and the same generation temperature for each gene. Comparing full precision against four-bit weights with a four-bit cache, quantization incurred only a modest change in mean amino-acid recovery, decreasing performance by 1.29 percentage points on average across the panel, with no significant difference between conditions (paired t-test, p = 0.23; Wilcoxon signed-rank test, p = 0.13; **Fig. 5C-D, Supplementary Table 3**). At the individual-gene level, *gyrA* showed a reproducible paired difference across the eight matched seeds (p = 0.0001), although the panel-wide comparison was not significant. No gene exhibited a catastrophic or deterministic collapse, and the observed per-gene differences were small relative to the variability across stochastic generations, particularly for genes that were already variable in full precision. Two genes also showed higher mean recovery under quantization, consistent with the stochastic variation of the task rather than a systematic improvement. Overall, these results indicate that four-bit quantization preserves Evo 2’s gene-completion performance on the original benchmark panel, with no evidence of a systematic loss in generative accuracy.

### Running Evo2 on a single GPU

Until now, using the largest configuration of Evo 2 meant securing a multi-GPU server. A researcher would have to fall back on the 7B model and accept its lower accuracy on variant interpretation and generation. The barrier is absolute rather than gradual: in bfloat16 the weights of Evo 2 40B occupy about 82 GB, so on an 80 GB card the model cannot be loaded, failing part-way through construction, at block 46 of 50, before a single base is read. Weight compression alone does not lift the barrier either, because the released quantizer compresses a model that is already resident and so presupposes the hardware that was missing. We resolve this by compressing each block as it is built and by distributing pre-quantized checkpoints, so the compressed model is reconstructed directly and the uncompressed weights are never materialised.

The practical consequence is that a single 80 GB GPU now runs the full model. Evo 2 40B occupies 33.8 GB of that card, peaks at 49.0 GB while scoring 32,768 bases, and sustains 883 tokens per second, leaving roughly 30 GB free. A researcher can score a 30-kilobase locus in about forty seconds, or sweep every single-nucleotide variant across a gene-sized window overnight, on one card and without a scheduler allocation for four. The compressed model is numerically the same model, the identical sequence scores a mean log-likelihood of -0.83931 both on one GPU from the compressed checkpoint and through the standard four-GPU path, agreeing to five decimal places. The download is smaller too, 33.8 GB against approximately 80 GB, so the cost of a first experiment falls as well.

To confirm that this is a usable capability and not merely a memory measurement, we repeated the BRCA1 saturation-genome-editing analysis described above end to end on a single card, starting from the distributed four-bit checkpoint. The model loads in nineteen seconds and is resident in 31.9 GiB. Scoring the same eight hundred variants requires 1,070 forward passes over 8,192-base windows, peaks at 42.5 GiB, leaves roughly thirty gigabytes free on an 80 GB card, and completes in two and a half hours. With four-bit weights and a four-bit cache, the model separates loss-of-function from functional and intermediate variants with an AUROC of 0.914 (95% confidence interval 0.892 to 0.934) and correlates with the experimental function score at a Spearman coefficient of 0.560, against 0.908 and 0.551 for the full-precision model across four GPUs. This configuration compresses the cache as well as the weights and so is not identical to the weight-only condition reported above; the three conditions span 0.906 to 0.914 in AUROC, a range consistent with the numerical variation expected from cache quantization and from differing reduction order under sharding, and small relative to the separation being measured. Those four GPUs complete the same analysis in twenty-eight minutes, so consolidating onto one card costs roughly fivefold in wall-clock time. The throughput advantage reported above is a property of single-token generation and does not extend to this scoring workload, which is dominated by prompt processing. What the single-card path buys is not speed but access: a clinically relevant variant sweep that cannot be loaded in full precision on one accelerator now runs overnight on a single GPU.

Generation behaves differently from the scoring workload above. In a single-token decode benchmark, the model reaches 883 tokens per second on one GPU compared with 641 tokens per second when sharded across four GPUs. This advantage is consistent with the communication overhead of moving activations between devices during autoregressive decoding. It does not extend to prompt-heavy workloads such as the BRCA1 analysis, where four-GPU execution is faster. For generation, however, consolidating the model onto one card therefore improves both accessibility and throughput.

The same benefit extends to the smaller configuration on a more modest card. On a single GPU held to a 40 GB memory budget, a capacity typical of the accelerators available on academic and cloud clusters, Evo 2 7B in full precision reaches 131,072 tokens and fails beyond it (**Fig. 3D**, **Table 1**). With four-bit weights and a four-bit cache the same budget reaches 1,048,576 tokens, the model’s full context, at a peak of 33.7 GB: an eightfold increase in reachable context on unchanged hardware, and a megabase of DNA held in a single context within forty gigabytes. Reaching the full million tokens requires the streaming prefill path, which reconstructs the cache one block at a time rather than in full; under the default fused-prefill path the same budget reaches 524,288 tokens, still a fourfold gain (Table 1). Between the two configurations the practical envelope is therefore straightforward: the 7B model spans a full megabase within a 40 GB budget, while the 40B model, which cannot be loaded in full precision on one 80 GB device at all, runs there compressed. The choice of model is substantially less constrained by available hardware.

### Long context inference

Evo 2 evaluates a sequence in one parallel pass only up to a fixed length, 65,536 tokens for the 40B configuration and 131,072 for the 7B; beyond that the sequence must be presented in chunks, with the model state carried across each boundary. Chunked prefill is therefore not an optimisation but the only tractable route to the long contexts Evo 2 is built for. When testing it, we found that it does not work. Scoring an 8,192-base human locus in chunks and comparing against the exact single-pass result, the per-base log-likelihoods were essentially uncorrelated with the ground truth (Pearson r = 0.003 to 0.297 depending on chunk size, against r = 1.000 for the unchunked control), and total log-likelihood differed by thousands of nats. The failure is silent: no error is raised, and the returned values sit in a plausible numeric range.

The cause is architectural. In the released implementation, a Hyena layer that already holds state routes to the single-token recurrent path, which begins by discarding all but the last position of its input. The resulting length-one tensor is then added to the residual stream and broadcast across the block, so that for every chunk after the first, all but one position is never processed by any Hyena layer. The eight attention layers continue to behave correctly, which is why the output remains superficially reasonable rather than collapsing. Feeding 256 tokens as four 64-token chunks returns 67 positions rather than 256, which exposes the mechanism directly. The repair follows from the structure of the operator. The Hyena recurrence is linear, so the state after a block of L tokens starting from a non-zero initial state decomposes exactly into the zero-state result, which the existing fast transform already computes, plus a term that propagates the incoming state forward. We implement this block-wise continuation, together with the corresponding correction to the block outputs and the short-filter history, and validate it at three levels: against single-token recursion in isolation, against a single parallel pass through the real layers, and end to end on both model configurations.

With the correction in place, chunked inference closely reproduces single-pass scoring to numerical round-off. Across chunk sizes from 512 to 8,192 on Evo 2 40B, Pearson correlation with the exact forward is 0.9998 or better and the total likelihood differs by under three nats over eight thousand bases, against r of approximately zero before the repair; the zero-boundary control remains bit-exact, confirming the comparison itself. Critically, the residual error does not accumulate with sequence length. On Evo 2 7B, whose single-pass ceiling is 131,072 tokens, scoring that full length in 128 chunks gives the same agreement as scoring it in eight (r = 0.99985 in both cases), so the number of boundaries crossed is not a source of drift. The same holds under compression: at the cache-compressed tier the correlation is 0.9984 or better, and at the fully compressed tier the residual is dominated by quantization rather than by chunking, since fifteen boundaries move the worst-case per-base error from 2.12 to 2.18. Two cautions follow. The 0.9998 figure above is the full-precision path; against an exact full-precision forward the fully compressed tier correlates at 0.992, the residual being quantization rather than chunking. And because that tier is what the long-context results run on, we measured it at the 40B’s full single-pass ceiling: thirty-one boundaries over 65,536 bases give r = 0.9918, matching the value at eight thousand bases and confirming that the chunking residual saturates rather than accumulating with boundary count.

The repair also determines what long-context inference costs in practice. Without it, the only numerically correct route past the single-pass ceiling is the released implementation’s own recipe for an oversized prompt: one parallel prefill of the first 65,536 tokens, then single-token advance for everything after. We measured that path rather than assuming a rate for it, timing decode latency against cache length on the full-precision 40B, from 55.8 ms at 1,024 positions to 63.2 ms at 65,536 and 122 ms at 524,288. Because latency grows with the cache, the cost of a traversal is the integral of that curve and not any single rate; quoting the terminal rate for the whole walk overstates it by roughly a third. Integrating the measured curve predicts a stepping walk that we then ran end to end to within 0.9 percent. On that basis, scoring 580,076 bases by single-token advance costs 13.7 hours, while the repaired chunked path completes the same sequence in 22.4 minutes at 432 tokens per second, a factor of about 36. The speed is owed to the block-wise continuation and not to compression: at equal context the full-precision chunked path is faster still, because four-bit weights must be reconstructed before each matrix multiply. What compression buys at length is reach rather than speed, since beyond roughly half a megabase the full-precision cache exhausts a four-GPU node and the comparison stops being one of time. This is what converts megabase-scale analysis from a calculation that must be planned around into one that can be run interactively, and it is obtained together with correctness rather than traded against it: the same block-wise continuation supplies both.

To confirm that this is a usable capability and not merely a throughput measurement, we scored a complete genome end to end in a single context. Mycoplasmoides genitalium G37 (NC_000908.2, 580,076 bases) is among the smallest genomes of any free-living organism, and its phylum shows the smallest four-bit perplexity penalty of any taxon in Supplementary Table 1, so the compressed model is a faithful instrument for it. The full genome was processed in one continuous pass in 22.4 minutes at 432 tokens per second, crossing 283 chunk boundaries and peaking at 179 GB across four GPUs, with no single device exceeding 45.3 GB, yielding a per-base likelihood for every position and a total genome log-likelihood of −542,436. The released implementation cannot perform this calculation in a single parallel pass at all, since its ceiling is 65,536 tokens, and its only correct alternative would take an estimated 13.7 hours. Reading an entire genome in one context is therefore a calculation that now completes while the user waits rather than one scheduled overnight.

## Discussion

TurboQuant-Bio shows that calibration-free compression transfers cleanly from its original setting, the attention caches of conventional transformers [12,13], to the weights and the hybrid architecture of Evo 2, and that it does so while preserving the biological behaviour Evo 2 is valued for. The four-bit configuration is near-lossless across perplexity spanning the tree of life, genomic classification, splice-site prediction, gene completion and clinically relevant variant-effect prediction, with small degradation across the evaluated benchmarks. Against the calibration-free round-to-nearest baseline, the advantage is specific to cross-taxa perplexity, where the four-bit degradation is smaller or more favourable on all fourteen taxa; on single-genome perplexity and on the four GUE tasks the two schemes are statistically indistinguishable at this sample size. Because the method needs no calibration data and no architectural change, it can be applied swiftly to a new model release, an advantage that matters in genomics, where curated in-domain data are scarce and organism-specific.

The most consequential result is access. In full precision the forty-billion-parameter model cannot be loaded onto a single 80 GB accelerator at all, its weights alone requiring 82.3 GB; compressed, it runs there, and on a four-GPU node it reaches its full one-million-token context in 175 GB of resident memory, where full precision exhausts the node beyond 524,288 tokens. On a single GPU constrained to a 40 GB memory budget, four-bit compression takes Evo 2 7B from 131,072 tokens to 1,048,576, an eightfold increase that places a full megabase of sequence within a single context on hardware already common in institutional clusters. The model’s capabilities are unchanged by any of this; what changes is the hardware required to use them.

These gains are not free. Reconstructing four-bit weights before each matrix multiplication makes prompt processing roughly half as fast as full precision at matched context, so compression is better understood as buying reach than speed: below the length at which full precision exhausts memory it is the slower option, and above that length there is no full-precision result to compare against. For single-token generation, where cost is dominated by reading weights from memory rather than by arithmetic, the fused kernels hold throughput within roughly a tenth of full precision (Fig. 4), so the memory savings are not paid for twice.

The correctness result carries a broader lesson. The chunked-prefill defect we identified had been present in released code, produced no error, and returned values in a plausible range; only comparison against an exact reference revealed it. Long-context genomic inference is exactly the regime in which such a reference is expensive to obtain and therefore rarely computed, so a silent failure can persist. We would encourage the field to treat agreement with an exact single-pass result as a routine acceptance test whenever a long-context path is used, and we release ours as one. Repairing the path also changes what is practical: scoring a complete 580-kilobase bacterial genome in a single context takes 22 minutes, against 13.7 hours for the token-by-token recurrence that is otherwise the only numerically correct route past the single-pass ceiling. A calculation that had to be planned around now completes while the user waits.

It is also worth noting that until this defect is repaired, the question of how much long-range context a genomic model actually exploits cannot be posed cleanly, because the measurement instrument itself was faulty; the repaired path makes that question answerable, and we regard it as an important direction rather than one we settle here. Several other directions remain. The cache-compression analysis here is specific to the hybrid architecture of Evo 2, in which only a minority of layers carries a cache, and it would be informative to characterize the same trade-offs on other genomic architectures. Adding strong calibration-based baselines [8,9] would sharpen the comparison at the lowest bit-widths, and extending the systematic evaluation to still longer contexts and to additional downstream assays would further map where compression is and is not safe for Evo 2. The gene-completion results also show variability across genes and, for some genes, across stochastic generations, supporting the reporting of per-gene results alongside aggregate accuracy.

Calibration-free compression reduces both the fixed cost of the weights and the growing cost of the attention cache with negligible loss of biological capability, bringing the largest open genomic model, at its full one-million-token context, within reach of a single compute node, and a megabase of context within reach of a single commodity GPU. By releasing the method as open, reproducible tooling, together with the acceptance test that exposed the chunked-prefill defect, we aim to lower the hardware barrier to working with Evo 2 across the tree of life.

## Methods

### Calibration-free weight quantization

TurboQuant-Bio compresses the weights of Evo 2 with a data-oblivious, rotation-based scalar quantizer following TurboQuant [12], one of a family of calibration-free geometric quantizers that also includes PolarQuant [13]. It requires no calibration data and no architectural change. The method rests on a geometric property of high-dimensional vectors: a random orthogonal rotation leaves a vector’s norm unchanged but drives the distribution of its normalized coordinates toward a fixed near-Gaussian target that is known a priori, so an optimal scalar quantizer, in the classical Lloyd-Max sense [19], can be designed once for that target and applied coordinate by coordinate with no observation of the data. It is important to be precise about scope, because the two things we compress use different mechanisms: only the weight quantizer uses the random rotation and its precomputed codebook; the attention cache, described below, is compressed by a simpler calibration-free integer scheme that applies no rotation. Both are free of calibration data, which is the property that matters for biology, but they are not the same method.

We quantize the weight matrix of every linear layer independently in blocks of 128 weights. For each block we store its length as a per-block scale, normalize the block, apply a fixed random rotation, and map each rotated coordinate to the nearest entry of a precomputed codebook, using closed-form levels at very low bit-widths and levels obtained by Lloyd’s algorithm [19] on the known target distribution otherwise. At inference the block is reconstructed on the fly by looking up the codebook entries, inverting the rotation, and rescaling. Only linear layers are quantized; embedding tables and normalization parameters are kept in full precision for stability. The one-time compression pass moves each weight matrix to host memory before quantizing it, so that the transient memory of compression itself is bounded and the procedure runs on the same hardware that will serve the model.

### Calibration-free attention-cache quantization

Evo 2 is built on the StripedHyena-2 architecture [2], which interleaves data-controlled convolutional operators, in the Hyena lineage [3], with a small number of layers of standard self-attention [20]. The convolutional layers maintain a fixed-size recurrent state rather than a growing cache, only the attention layers hold a cache that grows with sequence length, eight of fifty layers in the 40B configuration and five of thirty-two in the 7B configuration. The attention layers use grouped-query attention [21], in which several query heads share a single key/value head. Following established, calibration-free practice for attention caches [6,7], keys are quantized with a separate scale and zero-point for each channel, reduced over positions, while values are quantized with a separate scale and zero-point for each position, reduced over the feature axis. Both use asymmetric integer quantization at the chosen bit-width. We emphasize that this cache scheme applies no rotation and is distinct from the rotation-based weight quantizer described above; the two share only the property of needing no calibration data. Evo 2 applies rotary position embeddings [22] to keys and queries, and the cache stores the keys after this rotation. Because our cache quantizer is a simple scale-and-round map with no additional rotation, reconstruction and attention can be applied directly, with no inverse-rotation step to reconcile against the position embeddings.

The cache is held as a packed integer store: quantized codes at the chosen bit-width, with the most recent positions kept briefly in full precision and periodically folded into the packed store, so that the dominant long-context portion of the cache is always held at low precision while the memory used to fold in new positions stays bounded by the size of one chunk rather than the whole context. How cache compression behaves on a hybrid architecture of this kind, and how much memory it actually saves when the model is sharded across several accelerators, has to our knowledge not previously been examined; for protein language models, the recent TurboESM work adapted rotation-based cache quantization to the amino-acid activation manifold with a pipeline compatible with rotary position embeddings [15], a related but architecturally distinct setting from the hybrid convolutional-attention model studied here.

### Reference implementation for compressed long-context inference

Evo 2 uses the StripedHyena-2 architecture, with a growing KV cache only in its attention layers. We quantize the linear-layer weights and attention K/V only; activations and the fixed-size Hyena recurrent state remain uncompressed. Long sequences are processed by chunked prefill. The first chunk is evaluated in parallel, and subsequent chunks extend the existing inference state, with the attention and Hyena sequence offsets advanced after each call. This bounds transient prefill memory by the chunk size rather than the full context length.

Chunked prefill as released is not numerically valid. When a Hyena layer already holds inference state, the released dispatch routes multi-token input to the single-token recurrent path, which retains only the final position of its input; the length-one result is then broadcast over the residual stream, so every position but one in each subsequent chunk bypasses the Hyena operators entirely. We repair this with a block-wise continuation. Writing the modal recurrence as a linear map, the state after a block of L tokens starting from state s is the sum of the zero-state result, which the existing fast transform computes, and a term that carries s forward through the block; the block outputs receive the matching correction, and the short-filter path is corrected by prepending the retained history before convolution and discarding the corresponding leading outputs. The correction is applied automatically whenever a chunked context is used. We verify it at three levels: against single-token recursion for the isolated recurrence, against a single parallel pass through the real layers for each of the three Hyena variants, and end to end against exact single-pass scoring on both model configurations, at every compression tier and at context lengths up to the largest for which an exact reference can be computed.

Single-GPU deployment of the 40B configuration requires that the uncompressed weights are never resident. The released quantizer compresses a model that is already loaded, which presupposes hardware able to hold it. We instead intercept model construction and replace each block’s linear layers with their compressed form as soon as that block is built, so peak memory is the compressed model plus one uncompressed block rather than the whole uncompressed model, and we distribute pre-quantized checkpoints so that the compressed model can be reconstructed directly from disk. Loading from a pre-quantized checkpoint requires neither the original weights nor a multi-GPU machine.

The attention cache is stored as bit-packed segments with a small fp16 residual buffer for newly produced K/V. Keys use asymmetric per-channel quantization across positions, while values use asymmetric per-token quantization across features, with fp16 scales and zero-points. The residual is periodically quantized and appended as a new segment without rewriting the existing cache. This packed representation provides the reported memory savings; the separate fake-quantization path used in fidelity experiments quantizes and immediately dequantizes K/V and therefore does not reduce memory. To avoid reconstructing the complete cache, the reference implementation uses streaming attention, processing the packed cache one block at a time. Each block is dequantized, incorporated into an fp32 online-softmax accumulator, and released before the next block is read. Peak reconstruction memory therefore depends on the block size rather than the total context.

The reference weight path reconstructs the full weight matrix from its codebook indices, per-block scales and random rotation before applying an ordinary matrix multiplication. This cost is amortized during prefill but becomes dominant during single-token decoding, producing the large slowdown observed for unfused compressed generation. These reference paths establish the numerical baseline and expose the memory–latency trade-off addressed by the fused implementation.

### Fused kernels for compressed inference

We accelerate both weight and KV reconstruction with kernels selected automatically according to operation shape and bit-width. For weight operations with multiple activation rows, such as prefill, fused_linear_v2 caches the combined per-block scaling factors, moves the inverse rotation onto the smaller activation operand, dequantizes to bf16 and uses cuBLAS for the matrix multiplication. This is 2.8–6.5 times faster than the reference weight forward. For single-token decoding, the Triton-based [24] int4_gemv_decode reads the packed four-bit weights directly, applies the codebook, scales and activation rotation in the kernel, and accumulates in fp32 without materializing a bf16 weight matrix. This fused low-bit GEMV strategy is related in spirit to mixed-precision inference kernels such as MARLIN [25]. The fused implementation removes the approximately order-of-magnitude slowdown of the reference path and brings compressed-weight decoding close to the stock model’s throughput. For KV prefill, the first chunk has no cached prefix, so it is attended directly in bf16 using flash-backed scaled-dot-product attention and quantized only afterward for storage [23]. For single-token decoding, the fused int4 attention kernel reads packed K/V with coalesced accesses, dequantizes them in registers and incorporates them directly into the fp32 online-softmax calculation. Reconstructed K/V are never written to global memory. This gives an approximately sixfold speedup over the unfused streaming attention operation. The optimized int4 kernels are used for single-token decode, while prefill, cache-extension calls and other bit-widths use the corresponding batched or reference paths. All fused kernels are validated against the reference implementations, and multi-GPU launches use an explicit device guard so each kernel runs on the GPU containing the current sharded layer.

### Two deployment tiers and memory accounting

The two compressions can be combined or used separately, and we expose them as two tiers so that a user can match the trade-off to their hardware. The first tier compresses only the attention cache and keeps the weights in full precision. It reduces the memory cost that grows with context, keeps generation as fast as, or faster than, full precision, and reaches context lengths full precision cannot. It is the recommended default for generation. The second tier additionally compresses the weights for the smallest total footprint, at essentially no cost to generation speed thanks to the fused weight kernel described above, and is the right choice when fitting the model on limited hardware is the binding constraint.

Throughout, we distinguish two memory quantities. Resident memory is the persistent footprint, the weights plus the packed cache, which grows with context. Peak memory is the instantaneous maximum, which additionally includes transient buffers, principally a convolutional fast-transform buffer during the prompt pass and the reconstruction buffer during generation. These are governed by independent controls: the size of the chunk used to process a long prompt sets the transient prompt buffer, while the cache bit-width and streaming set the generation peak and the resident footprint. For the 40B model, which is sharded across accelerators, all memory figures are summed over the devices; single-device figures are reported for the 7B model. Memory is read from the accelerator’s allocator as maximum and current allocated bytes.

### Datasets, evaluation and reproducibility

We evaluate Evo 2 in its 40B and 7B configurations, using the StripedHyena-2 weights from the Arc Institute release [1]. Numerical faithfulness of compression is quantified by the mean absolute error between the outputs of the compressed and full-precision models on matched inputs. Genomic capability is assessed by perplexity on non-overlapping windows of the T4 bacteriophage genome, by cross-taxa perplexity on windows from fourteen taxa spanning viruses, bacteria, archaea and eukaryotes, by zero-shot classification on four GUE tasks [17], by splice-site prediction, by the Evo 2 gene-completion panel of twelve genes across six organisms [1], and by zero-shot variant-effect prediction on the BRCA1 saturation-mutagenesis assay [18], scoring reference against alternate alleles by a log-likelihood difference and reporting agreement with curated pathogenicity labels. Memory is measured as peak and resident accelerator memory, and generation speed as decode throughput over a fixed number of generated positions. The quantization baseline is round-to-nearest at matched bit-width; calibration-based methods are not evaluated here, since the comparison of interest is between calibration-free schemes [8,9]. All experiments were run on NVIDIA H100 nodes; single-GPU experiments used one H100, with the Evo 2 7B context sweep additionally constrained to a 40 GB memory budget to emulate a smaller accelerator. The random rotation is generated under a fixed seed and shared across layers and runs, so reported metrics are deterministic rather than averaged over random initializations. The complete pipeline and figure-generation scripts are released under the Apache 2.0 licence, matching both the upstream Evo 2 release and the distributed checkpoints.

## Data availability

Evo 2 model weights (40B and 7B) are available from the Arc Institute release [1]. The T4 bacteriophage genome is available from GenBank under accession AF158101. The GUE benchmark tasks [17] and the cross-taxa genomes used for perplexity (Supplementary Table 1) are publicly available. The BRCA1 saturation-genome-editing dataset is from Findlay et al. [18]. The Evo 2 gene-completion gene panel is described in the original Evo 2 publication [1].

## Code availability

TurboQuant-Bio is available at github.com/Georgakopoulos-Soares-lab/turboquant-bio under the Apache 2.0 licence, and the pre-quantized four-bit checkpoints for Evo 2 40B and Evo 2 7B are distributed at huggingface.co/michalakis99/turboquant-evo2-int4. Both are public and require no registration. The package requires an existing Evo 2 installation, which it deliberately does not install, since Evo 2’s dependencies are compiled against a specific CUDA version and accelerator. The interface is a single call that loads Evo 2 at a chosen tier with the long-context correction applied automatically, and a second that scores a sequence of any length; the compressed checkpoint is fetched and cached on first use. The repository ships the quantizer, the fused kernels, the block-wise continuation, the benchmarks, the figure-generation scripts and a three-level acceptance test whose numerical stage compares against recorded log-likelihoods rather than merely checking that the code runs.

## Acknowledgements

Research reported in this publication was supported by the National Institute of General Medical Sciences of the National Institutes of Health under award number R35GM155468.

## Author contributions

M.P., A.T., and I.G.S. jointly conceived the study. M.P. and A.T. designed the methodology, developed the software, performed the experiments and formal analyses, and generated the visualizations. M.P. and A.T. interpreted the results and wrote the original manuscript. I.G.S. provided supervision, contributed to interpretation of the results, and reviewed and edited the manuscript. I.G.S. acquired funding for the project. All authors reviewed and approved the final manuscript.

## Supplementary Information

**Supplementary Table 1 . Cross-taxa perplexity for Evo 2 40B (512-bp windows, N = 20 per taxon).** TQ = TurboQuant-Bio 4-bit; RTN = round-to-nearest int4. TurboQuant-Bio’s degradation was consistently smaller than with round-to-nearest.

Supplementary Table 2. Zero-shot BRCA1 variant-effect prediction with full-precision and four-bit Evo 2 40B.

Supplementary Table 3. Per-gene Evo 2 gene-completion performance with full-precision and four-bit weight and KV-cache configurations.

## References

[1] Brixi G, Durrant MG, Ku J, Naghipourfar M, Poli M, Sun G, et al. Genome modelling and design across all domains of life with Evo 2. Nature. 2026;652(8112):1349–1361. doi:10.1038/s41586-026-10176-5.

[2] Ku J, Nguyen E, Romero DW, Brixi G, Yang B, Vorontsov A, et al. Systems and algorithms for convolutional multi-hybrid language models at scale. arXiv:2503.01868; 2025.

[3] Poli M, Massaroli S, Nguyen E, Fu DY, Dao T, Baccus S, et al. Hyena Hierarchy: towards larger convolutional language models. International Conference on Machine Learning (ICML); 2023. arXiv:2302.10866.

[4] Nguyen E, Poli M, Durrant MG, Kang B, Katrekar D, Li DB, et al. Sequence modeling and design from molecular to genome scale with Evo. Science. 2024;386(6723):eado9336. doi:10.1126/science.ado9336.

[5] Kwon W, Li Z, Zhuang S, Sheng Y, Zheng L, Yu CH, et al. Efficient memory management for large language model serving with PagedAttention. ACM Symposium on Operating Systems Principles (SOSP); 2023. arXiv:2309.06180.

[6] Liu Z, Yuan J, Jin H, Zhong S, Xu Z, Braverman V, et al. KIVI: a tuning-free asymmetric 2-bit quantization for KV cache. International Conference on Machine Learning (ICML); 2024. arXiv:2402.02750.

[7] Hooper C, Kim S, Mohammadzadeh H, Mahoney MW, Shao YS, Keutzer K, Gholami A. KVQuant: towards 10 million context length LLM inference with KV cache quantization. Advances in Neural Information Processing Systems (NeurIPS); 2024. arXiv:2401.18079.

[8] Frantar E, Ashkboos S, Hoefler T, Alistarh D. GPTQ: accurate post-training quantization for generative pre-trained transformers. International Conference on Learning Representations (ICLR); 2023. arXiv:2210.17323.

[9] Lin J, Tang J, Tang H, Yang S, Chen WM, Wang WC, et al. AWQ: Activation-aware Weight Quantization for On-Device LLM Compression and Acceleration. Proceedings of Machine Learning and Systems. 2024.

[10] Ashkboos S, Mohtashami A, Croci ML, Li B, Cameron P, Jaggi M, et al. QuaRot: outlier-free 4-bit inference in rotated LLMs. Advances in Neural Information Processing Systems (NeurIPS); 2024. arXiv:2404.00456.

[11] Liu Z, Zhao C, Fedorov I, Soran B, Choudhary D, Krishnamoorthi R, et al. SpinQuant: LLM Quantization with Learned Rotations. International Conference on Learning Representations (ICLR); 2025. arXiv:2405.16406.

[12] Zandieh A, Daliri M, Hadian M, Mirrokni V. TurboQuant: online vector quantization with near-optimal distortion rate. International Conference on Learning Representations (ICLR); 2026. arXiv:2504.19874.

[13] Han I, Kacham P, Karbasi A, Mirrokni V, Zandieh A. PolarQuant: quantizing KV caches with polar transformation. International Conference on Artificial Intelligence and Statistics (AISTATS); 2026. arXiv:2502.02617.

[14] Tseng A, Chee J, Sun Q, Kuleshov V, De Sa C. QuIP#: even better LLM quantization with Hadamard incoherence and lattice codebooks. International Conference on Machine Learning (ICML); 2024. arXiv:2402.04396.

[15] Hu Y, Wang J, Liu Y. TurboESM: ultra-efficient 3-bit KV cache quantization for protein language models with orthogonal rotation and QJL correction. arXiv:2603.26110; 2026.

[16] Avsec Ž, Agarwal V, Visentin D, Ledsam JR, Grabska-Barwinska A, Taylor KR, et al. Effective gene expression prediction from sequence by integrating long-range interactions. Nature Methods. 2021;18(10):1196–1203. doi:10.1038/s41592-021-01252-x.

[17] Zhou Z, Ji Y, Li W, Dutta P, Davuluri R, Liu H. DNABERT-2: efficient foundation model and benchmark for multi-species genomes. International Conference on Learning Representations (ICLR); 2024. arXiv:2306.15006.

[18] Findlay GM, Daza RM, Martin B, Zhang MD, Leith AP, Gasperini M, et al. Accurate classification of BRCA1 variants with saturation genome editing. Nature. 2018;562(7726):217–222. doi:10.1038/s41586-018-0461-z.

[19] Lloyd SP. Least squares quantization in PCM. IEEE Transactions on Information Theory. 1982;28(2):129–137. doi:10.1109/TIT.1982.1056489.

[20] Vaswani A, Shazeer N, Parmar N, Uszkoreit J, Jones L, Gomez AN, et al. Attention is all you need. Advances in Neural Information Processing Systems (NeurIPS); 2017. arXiv:1706.03762.

[21] Ainslie J, Lee-Thorp J, de Jong M, Zemlyanskiy Y, Lebrón F, Sanghai S. GQA: training generalized multi-query transformer models from multi-head checkpoints. Conference on Empirical Methods in Natural Language Processing (EMNLP); 2023. arXiv:2305.13245.

[22] Su J, Lu Y, Pan S, Murtadha A, Wen B, Liu Y. RoFormer: enhanced transformer with rotary position embedding. Neurocomputing. 2024;568:127063. arXiv:2104.09864.

[23] Dao T, Fu DY, Ermon S, Rudra A, Ré C. FlashAttention: fast and memory-efficient exact attention with IO-awareness. Advances in Neural Information Processing Systems (NeurIPS); 2022. arXiv:2205.14135.

[24] Tillet P, Kung HT, Cox D. Triton: an intermediate language and compiler for tiled neural network computations. ACM SIGPLAN International Workshop on Machine Learning and Programming Languages (MAPL); 2019. doi:10.1145/3315508.3329973.

[25] Frantar E, Castro RL, Chen J, Hoefler T, Alistarh D. MARLIN: mixed-precision auto-regressive parallel inference on large language models. arXiv:2408.11743; 2024.

